# Cortical interneuron development is affected in leukodystrophy 4H

**DOI:** 10.1101/2022.08.22.504736

**Authors:** Stephanie Dooves, Liza M.L. Kok, Dwayne B. Holmes, Nicole Breeuwsma, Marjolein Breur, Marianna Bugiani, Nicole I. Wolf, Vivi M. Heine

## Abstract

4H leukodystrophy is a rare genetic disorder classically characterized by hypomyelination, hypodontia and hypogonadotropic hypogonadism. With the discovery that 4H is caused by mutations that affect RNA polymerase III, mainly involved in the transcription of small non-coding RNAs, also patients with atypical presentations with mainly a neuronal phenotype were identified. Pathomechanisms of 4H brain abnormalities are still unknown and research is hampered by a lack of preclinical models. We aimed to identify cells and pathways that are affected by 4H mutations using induced pluripotent stem cell models.

RNA sequencing analysis on induced pluripotent stem cell-derived cerebellar cells revealed several differentially expressed genes between 4H patients and control samples, including reduced *ARX* expression. As *ARX* is involved in early brain and interneuron development, we studied and confirmed interneuron changes in primary tissue of 4H patients. Subsequently, we studied interneuron changes in more depth and analyzed induced pluripotent stem cell-derived cortical neuron cultures for changes in neuronal morphology, synaptic balance, network activity and myelination. We showed a decreased percentage of GABAergic synapses in 4H, which correlated to increased neuronal network activity. Treatment of cultures with GABA antagonists led to a significant increase in neuronal network activity in control cells but not in 4H cells, also pointing to lack of inhibitory activity in 4H. Myelination and oligodendrocyte maturation in cultures with 4H neurons was normal, and treatment with sonic hedgehog agonist SAG did not improve 4H related neuronal phenotypes. qPCR analysis revealed increased expression of parvalbumin interneuron marker *ERBB4*, suggesting that the development rather than generation of interneurons may be affected in 4H.

Together, these results indicate that interneurons are involved, possibly parvalbumin interneurons, in disease mechanisms of 4H leukodystrophy.

## Introduction

Leukodystrophies are genetic disorders characterized by primary brain white matter involvement. In children, leukodystrophies are often progressive and can lead to early death.(1) Until several years ago, up to half of patients with leukodystrophies did not receive a genetic diagnosis. Since, developments in next generation sequencing techniques have led to the rapid identification of gene mutations underlying different childhood leukodystrophies, such as *de novo* mutations in a structural genes, e.g. *TUBB4A*, in hypomyelination with atrophy of the basal ganglia and cerebellum (OMIM 612438) or mutations in genes encoding proteins essential for translation, e.g. *DARS* (OMIM 615281) or *RARS* (OMIM 616140) in hypomyelinating leukodystrophies(2-5). Insights into genetic causes allowed incredible progress in diagnostics, but our understanding of the mechanisms responsible for disease pathology is still lacking(6, 7).

One of the more prevalent leukodystrophies is 4H syndrome (OMIM 612440), originally characterized by hypomyelination, hypogonadotropic hypogonadism, and hypodontia(8-10). Other characteristics are cerebellar atrophy and myopia. Epilepsy has been described in some patients. Whole exome sequencing has revealed that 4H leukodystrophy is caused by abnormal RNA polymerase III (POLR3), and so far variants in genes encoding different POLR3 subunits, *POLR3A, POLR3B, POLR1C* and *POLR3K*, have been identified(11-14). POLR3 is responsible for the transcription of many different classes of non-protein coding (nc) RNAs, with diverse biological functions, such as transfer (t), ribosomal (r), small nuclear (sn), small nucleolar (sno), and micro (mi) RNAs(15). Given the variety of 4H presentations and the diverse regulatory functions of POLR3 genes, there are many potential pathways and tissue-types to research. Interestingly, some POLR3 mutations do not lead to classic brain white matter defects associated with 4H, but rather show a predominant neuronal phenotype with involvement of the basal ganglia(10, 16, 17). So while 4H was originally described as a typical hypomyelinating disorder, it suggests that POLR3 variants could also primarily affect neuronal populations.

The generation of induced pluripotent stem cells (iPSCs) from patient tissue allows for patient-specific disease models starting from early embryonic stage(18), and therefore provide a promising tool to study rare diseases like 4H leukodystrophy. To get insight into disease mechanisms and identify genes that are dysregulated by 4H mutations, we generated patient iPSC-derived cerebellar and cortical cell populations, including neuron-oligodendrocyte co-cultures and studied iPSC products by whole transcriptome, cellomics and multi-electrode array (MEA) analysis. In patient iPSC-derived cerebellar neurons we found a significantly lower expression of the Aristaless Related Homeobox (*ARX*) gene, which is involved in early brain development and severe infantile-onset encephalopathies(19). As ARX is associated with the development of cortical interneurons, we performed immunohistochemistry for GAD65/67 on patient material and confirmed alterations in populations of inhibitory neurons in 4H. So we decided to study the development of cortical neurons in more depth. Indeed, the number of GABAergic synapses was significantly decreased in 4H neuronal cultures, which correlated to an increased neuronal network activity. To further identify the specific subtypes that are affected in 4H, qPCR analysis for the major cortical interneuron populations was performed. Interestingly, the expression of *ERBB4*, important for the development of parvalbumin interneurons, was significantly increased in 4H neurons. Together, our results show that cortical interneuron development is affected in 4H patients, possibly due to pathway changes involving ARX that may affect development of parvalbumin interneurons.

## Material and methods

### iPSC culture

Fibroblasts from anonymous donors and 4H patients were reprogrammed into iPSC lines as described previously for controls, with written informed consent of patients/guardians(20). Briefly, iPSCs were generated using an overexpression of *OCT4, SOX2, KLF4*, and *C-MYC*. iPSC lines were confirmed for pluripotency by ICC, RTPCR, alkaline phosphatase, embryoid body formation assay, karyotyping or a pluritest. Human ESC control line H01 was obtained from WiCELL and used as a control line for RNA sequencing experiments. See Table 1 for an overview of the iPSC lines. Control and 4H iPSCs were maintained on a vitronectin coating in TeSR E8 medium. Medium was refreshed daily, and cells were passaged once a week using Gentle Cell Dissociation reagent (StemCell Technologies) according to manufacturer’s protocol. Cells were split 1:10-1:50 to a new well for further maintenance or start of differentiation protocols.

**Table 1.** iPSC lines.

| **Line** | **Genotype** | **Age** | **Sex** | **Reprogramming method** | **Used for experiments** |
| --- | --- | --- | --- | --- | --- |
| C1 | Control | 3d | M | Lentiviral | RNA sequencing |
| C2 | Control | 74d | M | Lentiviral | All |
| C3 | Control | 70y | M | Lentiviral | RNA sequencing |
| C4 | Control | 19y | F | Lentiviral | Neuronal analysis, Neuron-glia co-culture, qPCR |
| C5 | Control | 21y | M | Lentiviral | All except RNA sequencing |
| C6 | Control | 19y | M | Lentiviral | All except RNA sequencing |
| P1 | 4H<br>(POLR3B) | 10y | F | Lentiviral | All |
| P2 | 4H<br>(POLR3B) | 2y | M | Lentiviral | All |
| P3 | 4H<br>(POLR3B) | 4.5y | F | Lentiviral | All |
| P4 | 4H<br>(POLR3A) | 12y | F | Lentiviral | All except RNA sequencing |
| P5 | 4H<br>(POLR3A) | 24y | M | Sendai | qPCR interneuron subtypes |
| P6 | 4H<br>(POLR3A) | 2y | F | Sendai | qPCR interneuron subtypes |

### Differentiation in neural cells

Human PSCs were kept in feeder-free, serum-free cell culture as described previously(20), and then differentiated into neural cell types. Differentiation into cerebellar granule cell neurons was done using a previously reported protocol, involving minimal factors in an N2+B27 neural maintenance medium (NMM), modified for a two-dimensional (2D) monolayer culture(20, 21). Cortical neuron differentiation was performed as previously described(22) using dual SMAD inhibition and consequent maturation of cells into both glutamatergic and GABAergic neurons. For myelinating cultures, human iPSCs were differentiated into glial progenitor cells as previously described(23). See supplementary method section for more details on culture protocols.

### Neuron-glia co-culture

For myelinating neuron-glia cultures, neurons and glia cells were differentiated from iPSCs as described in the supplementary methods. To promote neuronal maturation, neurons were plated on a monolayer of rat astrocytes. Rat astrocytes were isolated from postnatal day 1 rat cortex using papaïn dissociation. At 1 day and 4 days after dissociation, flasks with rat astrocytes were tapped to detach contaminating cells like oligodendrocytes or microglia as much as possible. Astrocytes were frozen at passage 1 until further use. To start a co-culture, day 18 cortical neurons were plated on a monolayer of rat astrocytes in Neurobasal/B27 medium supplemented with BDNF, GDNF, cAMP and IGF1. At day 30 of neuronal differentiation, day 67 human glial progenitor cells were added to the culture, and medium was switched to Neurobasal/N1 medium supplemented with T3, NT3, mouse Laminin, BDNF and cAMP. The co-cultures were kept in culture for an additional 40 days, at which point cultures were fixed for immunostaining.

### RNA sequencing

Total RNA was isolated from cells using TRIzol and chloroform-isopropanol extraction. After quality control samples were run on an Illumina HiSeq2500 according to manufacturer’s protocol. Sequencing reads were aligned to Humane Genome hg38. Similarity between fibroblast, iPSC and cerebellar cells was analyzed, and the data of iPSC and cerebellar cells were compared to previously published gene expression profiles of 2 iPS cells and 2 granule cells. Differentially expressed genes between 4H and control were determined with DESeq2 package in R. See supplementary methods for more details.

### qPCR

For qPCR analysis RNA was isolated from neurons using TRIzol-chloroform isolation. After production of cDNA, a qPCR was performed using SYBR green according to manufacturer’s protocol. For *ARX* qPCR data was normalized for housekeeping gene *EIF4G2*. For interneuron subtype analysis, data was normalized for housekeeping gene *EIF4G2* and general neuronal marker *NEUN*. See Supplementary Table 1 for an overview of the primers used. From the results the fold change 2^-δδ^^CT^ values were calculated, and the log fold change was used for statistical analysis.

### Immunocytochemistry

Cells were fixed for immunostaining using either 4% PFA or methanol:acetone fixation. Primary antibodies (Supplementary Table 2) were incubated overnight, followed by 2 hour incubation with secondary antibody and embedding with Fluoromount G. See supplementary methods for more details.

### Immunohistochemistry

Cortical tissue was obtained from a control (F, 7mo, cause of death: hypoxic ischemic encephalopathy) and a 4H patient (F, 16y). The tissue was formalin-fixed paraffin-embedded (FFPE) and 5μm-thick sections were made. Immunofluorescence staining was started by rehydrating sections in xylene and alcohol. Endogenous peroxidase was quenched in 0.3% (w/v) H_2_0_2_ in PBS for 30 minutes, followed by heat induced antigen retrieval in citrate buffer (pH=6). Primary antibody GAD65/67 (1:1000, G-5163, Sigma) was incubated overnight at room temperature. The following day slides were rinsed and incubated with horseradish peroxidase labelled secondary antibodies for 1 hour and developed using 3,3′-diaminobenzidine (DAB, 1:50, DAKO) for 5 minutes. Sections were counterstained with haematoxylin, dehydrated with alcohol and xylene and mounted with Quick-D (Klinipath, 7280). Light microscopy pictures were taken with a Leica DM6000B microscope (Leica microsystems).

### SAG treatment

SAG (Cayman Chemical) was dissolved in DMSO to a stock concentration of 100 μM. In a pilot experiment 1 line of control and 1 line of 4H neurons were treated with 3 nM, 50 nM, 100 nM or 250 nM SAG either starting from day 18 or day 30 on. Based on the pilot experiment a concentration of 100 nM SAG was chosen, with treatment starting from day 18 until the endpoint of the experiment. SAG or vehicle (1:1000 DMSO) was added twice a week with the medium refreshment.

### Multi electrode array

Neuronal activity was measured on 24 well multi-electrode array (MEA) plates (Multi Channel Systems 24W300/30G-288). Cultures were measured weekly for 30 minutes, after which data was analyzed with Multiwell-Analyzer software (Multichannel Systems) to determine spike count, burst count and network burst counts from raw data. See supplementary methods for more details.

### Statistical analysis

All experiments were repeated on at least 2-3 independent cell batches per patient, and the results of those experiments were averaged to obtain one value per iPSC line. MEA data was collected on 1 batch of neurons per cell line. All analyses were either done automated or blinded. Neuronal morphology, myelination and the number of OLIG2 and MBP positive cells were analyzed using Columbus software (Perkin Elmer). Synaptic analyses and the percentage of myelinated axons were determined in ImageJ (imagej.nih.gov/ij) using the NeuronJ and SynaptoCount plugins (see Supplementary Methods). Statistical analyses were done in IBM SPSS statistics software version 26 (IBM). Data was tested for normality with a Shapiro-Wilk test. Significance was tested with independent samples t-tests (for parametric data) or a Mann-Whitney U test (for non-parametric data). To compare the effect of compounds on MEA data or the effect of SAG treatment paired samples t-test or Wilcoxon signed rank tests were used. All statistical tests were two-tailed and tested against alpha-level *P* < 0.05.

## Data availability

The authors confirm that the data supporting the findings of this study are available within the article and its supplementary material.

## Results

### *In vitro* products present with iPSC and cerebellar cell identity

As 4H patients with *POLR3B* variants often have cerebellar atrophy, we reprogrammed patient fibroblasts into iPSCs and differentiated these into cerebellar neurons as described previously(21). We first wanted to establish that reprogramming and differentiation products derived from patient and control fibroblasts were unique cell populations, with relatively little variation due to source material. Further, we tested the validity of the reprogramming protocols by estimating how close each derived cell type resembled published expression profiles. To investigate sample similarity, t-Distributed Stochastic Neighborhood Embedding (t-SNE) was performed with perplexity at 5. Cluster analysis showed that samples group together based on cell types (fibroblast, iPSC, product) regardless of cell source (patient or control) (Fig. 1A). The relatively high similarity of samples from the same cell type, regardless of their source, suggests that reprogramming methods were effective at generating a consistent cell type.

**Figure 1.**
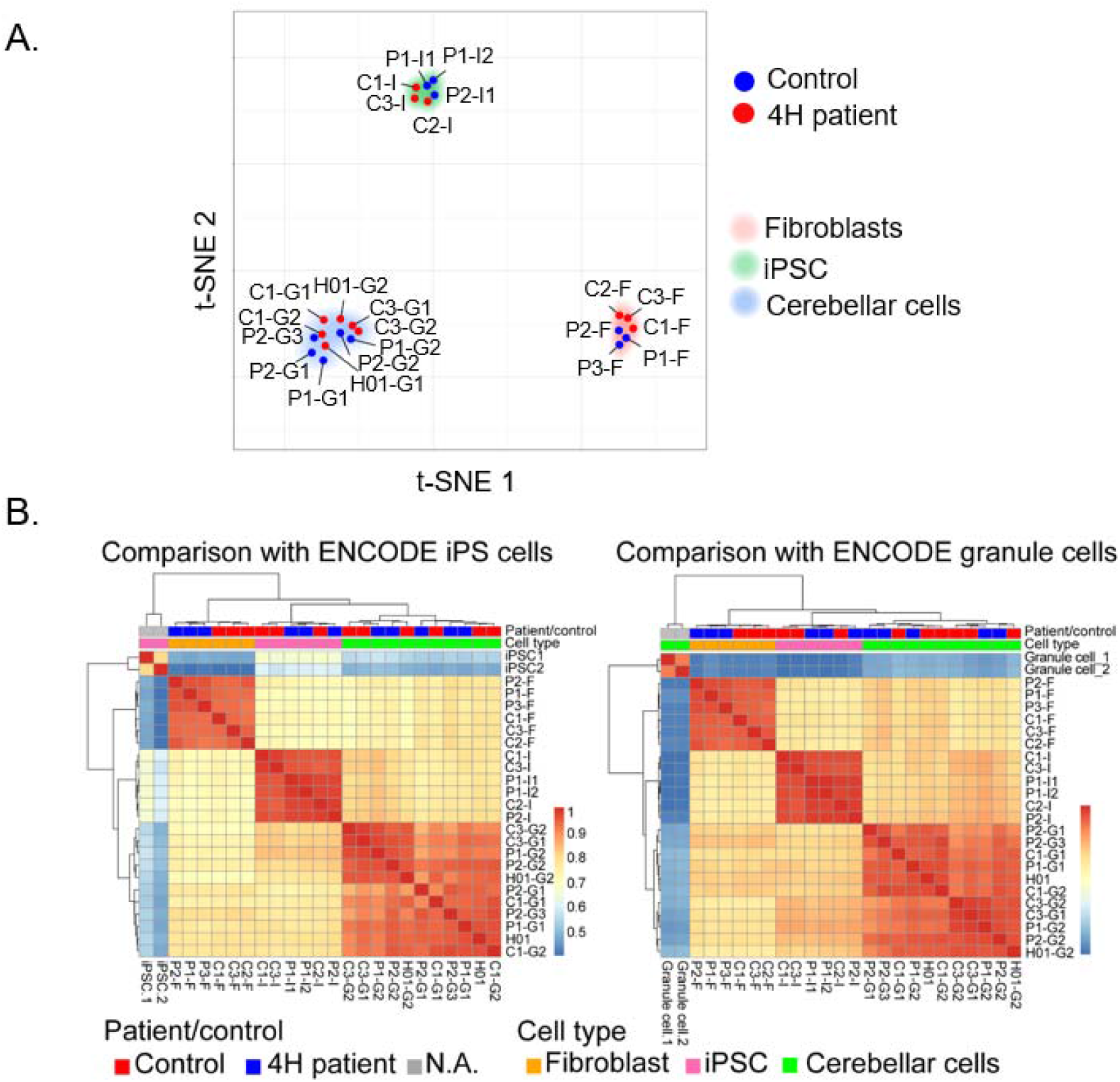
Reprogramming and differentiation leads to distinct cell types. (**A**) shows the projection of 23 samples in 2-dimensional space. Each dot represents a sample that is colored by patient/control status. Samples are positioned based on relative similarity, showing that fibroblasts, iPSCs and cerebellar cells cluster together. (**B**) shows heatmaps of the comparison of expression profiles with ENCODE iPSC and granule cell samples. Heatmap is colored by spearman’s rank correlation coefficient. Patient/control status and cell types are colored at the top of the heatmap. (**A**-**B**) P = patient, C = control, I = iPSC, F = fibroblast, G = cerebellar granule cell.

To estimate the accuracy of cell reprogramming, pair-wise Spearman’s rank correlations were computed for 23 samples from this study and two expression profiles obtained from ENCODE. One was an ENCODE iPSC expression profile (ENCSR722POQ) and the other an ENCODE granule cell expression profile (ENCSR313IUO), both containing two biological replicates (Fig. 1B). Hierarchical clustering showed that samples in both analyses group by cell type. Color coding indicates that iPSCs and cerebellar cells are more closely related to the relevant ENCODE profiles, than the other cell types (iPSCs = *P* < 0.001; cerebellar cells = *P* < 0.001). Taken together, fibroblast-derived iPSCs and PSC-derived cerebellar cells can be treated as relevant representatives of those developmental stages/lineages for further analyses.

### Transcriptome analysis revealed 4H-associated changes in gene expression

RNA sequencing analysis showed several differentially expressed genes (DEGs) between 4H patients and controls. In fibroblasts, 29 genes showed a significantly altered expression (Supplementary Table 3), while in iPSCs 20 genes were differentially regulated (Fig. 2A, Supplementary Table 4). Notable DEGs that were downregulated in 4H iPSCs include several genes involved in neural development (*OTX2, NPTX1, SLITRK6, SEMA6A*) and a downregulation of *TMEM64* that plays a role in osteoclast differentiation(24-28).

**Figure 2.**
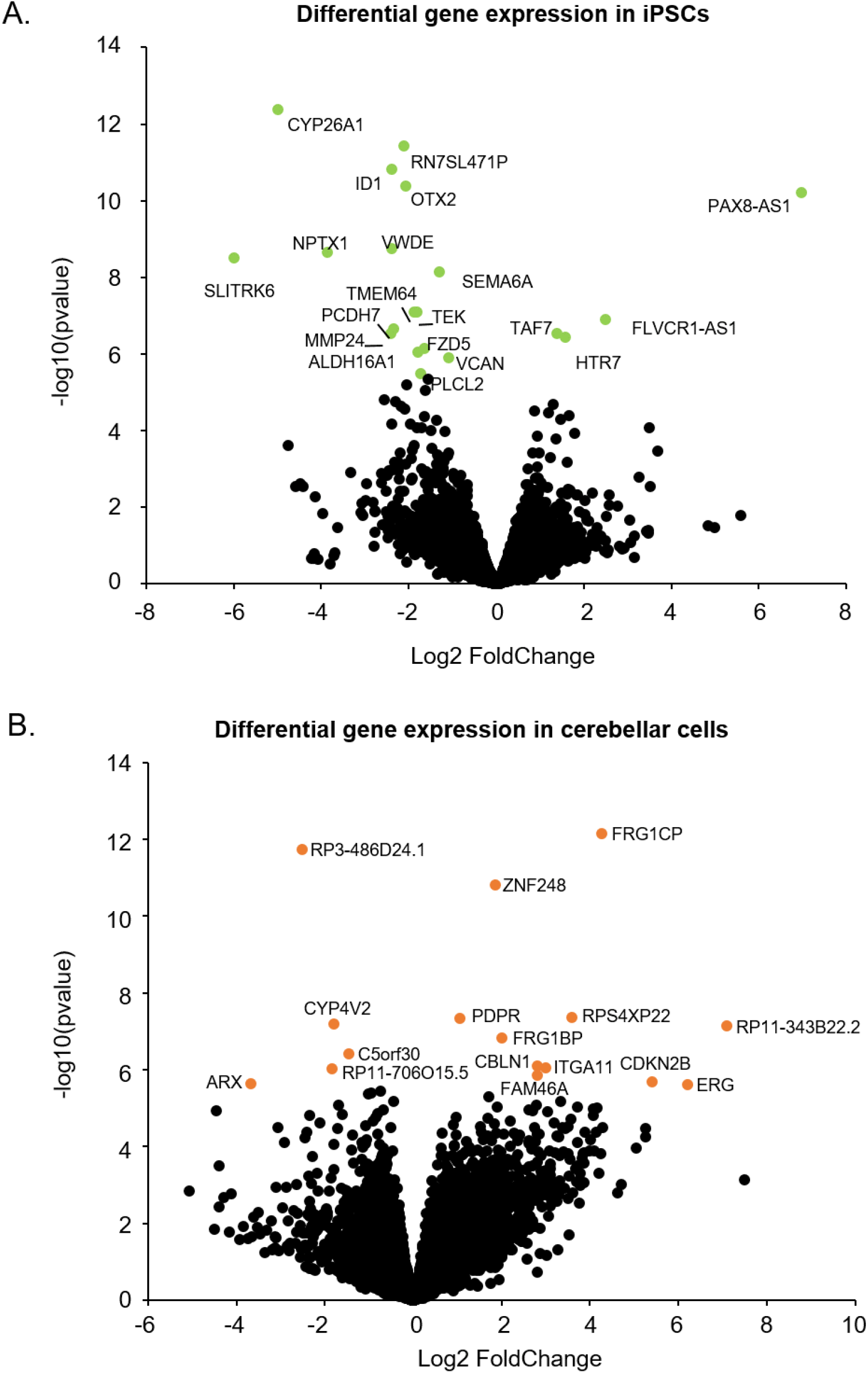
Differential gene expression reveals genes involved in neurodevelopment. (**A**) shows a volcano plot of the RNA sequencing data on patient and control iPSCs. DEGs are visualized by a green dot and gene label. (**B**) Volcano plot showing DEGs in 4H cerebellar cells. The significant DEGs are indicated with an orange dot and gene label.

In 4H cerebellar cells, 16 genes showed a significant differential expression (Fig. 2B, Supplementary Table 5). 4H cerebellar cells showed an increased expression of *PDPR*, which is associated with intellectual disability(29) and an increased expression of *CBLN1*, a cerebellum-specific precursor protein(30). The downregulation of *ARX* appears to be a potentially significant discovery. *ARX* mutations are associated with several brain disorders, and *ARX* loss of function is associated with abnormal cortical interneuron development and migration(31-34).

### Cortical interneuron changes in 4H

We studied whether the decreased *ARX* expression point to cortical interneuron involvement in 4H. iPSCs were differentiated into cortical neurons, i.e. a mixture of GABAergic interneurons and glutamatergic cortical projection neurons, according to previously established protocols(22). At a neuronal precursor cell state, cells were harvested for RNA analysis. Consistent with the cerebellar cells, *ARX* expression was decreased in immature neurons of 4H patients (control 1.08 ± 0.31, 4H 0.26 ± 0.10, *t*(6) = 12.073, *P* < 0.001, Fig. 3A-B), confirming that *ARX* dysregulation may be a common feature in 4H.

**Figure 3.**
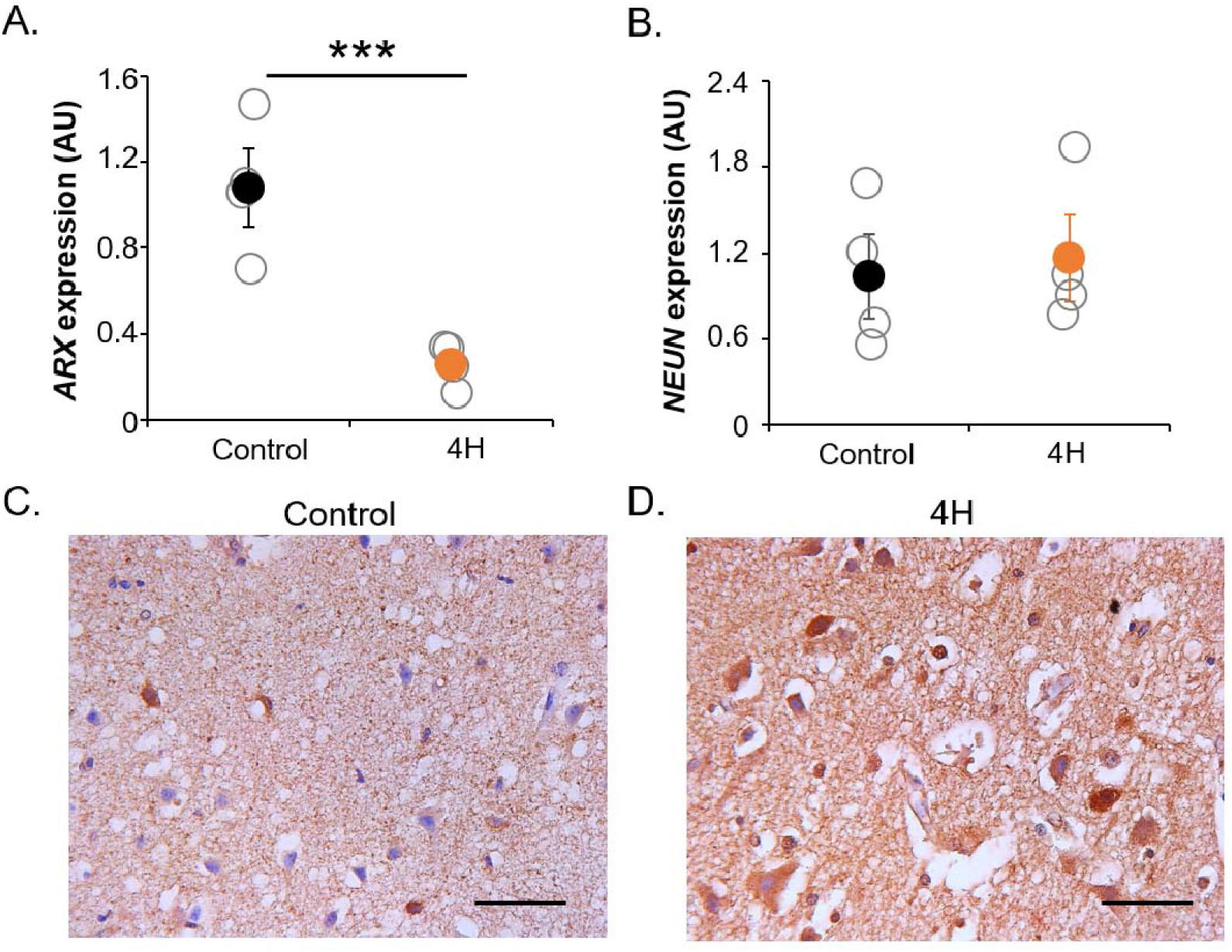
Cortical interneuron changes in 4H. (**A**) QPCR analysis shows ARX expression in 4H and control neurons. RNA was isolated from cortical neurons at day 18 of the differentiation protocol, which represents an immature state. In 4H neurons, ARX expression is significantly downregulated, while there is no change in the expression of pan-neuronal gene NEUN (**B**). *** = P < 0.001, AU = arbitrary units. Open circles represent average per individual patient/control, solid circles represent the mean per genotype. Error bars represent standard error of the mean. Immunohistochemical analysis for GAD65/67 on primary cortical tissue of a control individual (**C**) and 4H patient (**D**) showed that the pan interneuronal marker is increased in 4H brain tissue. Scale bar =100 µm.

To confirm that cortical interneurons are affected in primary brain tissue of 4H patients, we studied patient tissue for expression of GAD65/67, a pan GABAergic marker widely used to visualize all inhibitory neurons. Patient tissue showed elevated immunoreactivity for GAD65/67 compared to control, strengthening the hypothesis that cortical interneuron populations are affected in 4H (Fig. 3C-D). As patient tissue is really scarce and iPSC-based model systems present differential gene expression in 4H cells, we continued functional analysis in iPSC-derived cortical cultures.

### 4H neurons show a decreased generation of GABAergic synapses

To investigate how cortical neuron development is affected in 4H, cortical neuronal cultures were matured and analyzed for morphology changes (Fig. 4). Morphology of 4H and control neurons was analyzed on dendritic (MAP2) and axonal (NF-200) staining using different parameters, including axonal and dendritic length, branching level, and number of extremities and roots. No significant change in any parameter for neuronal morphology was observed between 4H and control neurons (Fig. 4B-C).

**Figure 4.**
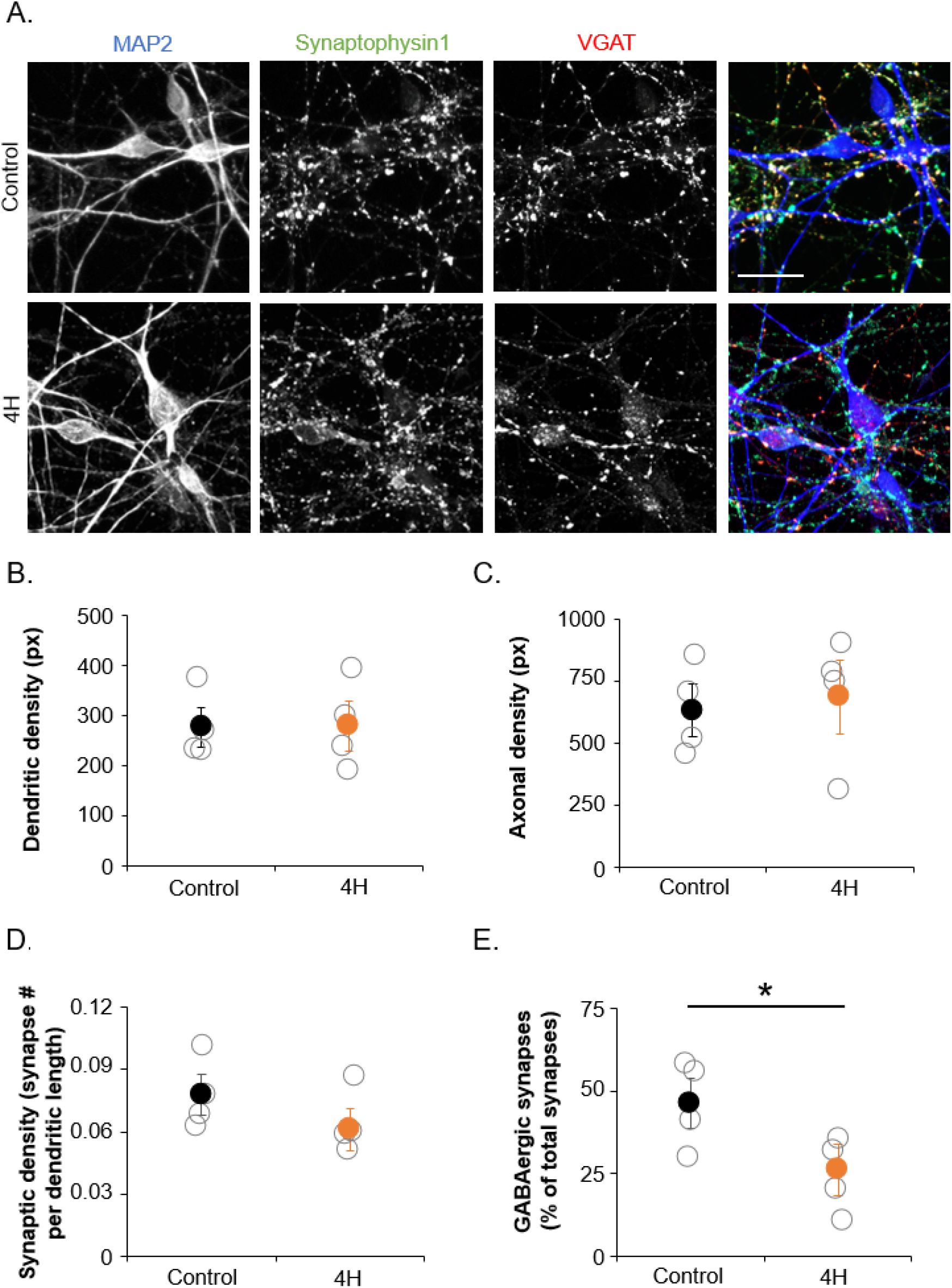
Cortical neurons of 4H patients show a decreased generation of GABAergic synapses. (**A**) shows representative images of immunostainings on control and 4H neurons. Neurons are stained for dendrites (MAP2, blue), all synapses (Synaptophysin1, green) and GABAergic synapses (VGAT, red). Dendritic (**B**) and axonal (**C**) density were not changed in 4H neurons compared to controls. The number of synapses per dendritic length was normal in 4H (**D**), while the percentage of GABAergic synapses was significantly decreased in 4H neurons (**E**). Scalebar = 25 μm, px = pixel, * = P < 0.05. (**B**-**E**) open circles represent average per individual patient/control, solid circles represent the mean per genotype. Error bars represent standard error of the mean.

To analyze whether 4H cultures present changes in the synapses, we analyzed synaptic marker expression by immunocytochemistry (Fig. 4A). In our cortical neuronal cultures cells differentiate into GABAergic interneurons and glutamatergic neurons(22), allowing us to study differentiation efficiency into these neuronal subtypes. At day 56 of differentiation, we studied the total number of synapses by quantifying the number of synaptophysin1 puncta per MAP2 dendritic length (Fig. 4D). We observed no significant change between the 4H and control cultures (control 0.078 ± 0.017, 4H 0.061 ± 0.018, Fig. 4D). To study whether the fraction of inhibitory synapses changed in 4H cultures, we measured the number of GABAergic presynaptic terminals by co-localization of pre-synaptic protein synaptophysin1 puncta with vesicular GABA transporter (VGAT) puncta (Fig. 4E). While 4H neurons showed a normal amount of synapses, the percentage of GABAergic synapses was significantly decreased from 46.4% in control to 26.1% in 4H neurons (*t*(6) = 2.503, *P* = 0.046, Fig. 4E). The decreased generation of GABAergic synapses confirms that interneuron development is affected in 4H.

### 4H neurons show altered network activity

To assess whether the decreased percentage of GABAergic synapses also had consequences for network activity, we recorded spontaneous activity of developing neuronal networks from 3 controls and 4 4H patients plated on multi-electrode arrays (MEAs) (Fig. 5A). The activity was captured in three parameters namely single spike events, local bursts (spike trains) and network wide bursting events (Fig. 5B). Spontaneous action potential firing in the first weeks mainly consisted of single spike events, while local and network wide bursts occurred as the cultures matured (Fig. 5D-F). Tetrodotoxin administration (TTX, 1 uM) confirmed that baseline activity was depending on sodium ion channels (Supplemental Fig. 1A,D). At 14 weeks post plating, 3 out of 4 4H patients showed high levels of local and network bursts, which was mostly absent in control cultures (Fig. 5C,F). This is an indication of hyperactivity in 4H networks, although the network activity was not significantly different from control (spikes control 1.19 ± 0.40, 4H 2.41 ± 1.88; bursts control 1.65 ± 0.13, 4H 5.61 ± 5.21; network bursts control 0.33 ± 0.60, 4H 5.17 ± 5.16). Interestingly, the single 4H line (P3) that did not show an increased number of bursts and network bursts was also the 4H line with the highest percentage of GABAergic synapses. A correlation analysis could indeed confirm significant correlations between the percentage of VGAT^+^ synapses and the number of bursts (*r* = −0.774, *P* = 0.041) and network bursts (*r* = −0.786, *P* = 0.036), showing a relation between the amount of GABAergic pre-synapses and network activity.

**Figure 5.**
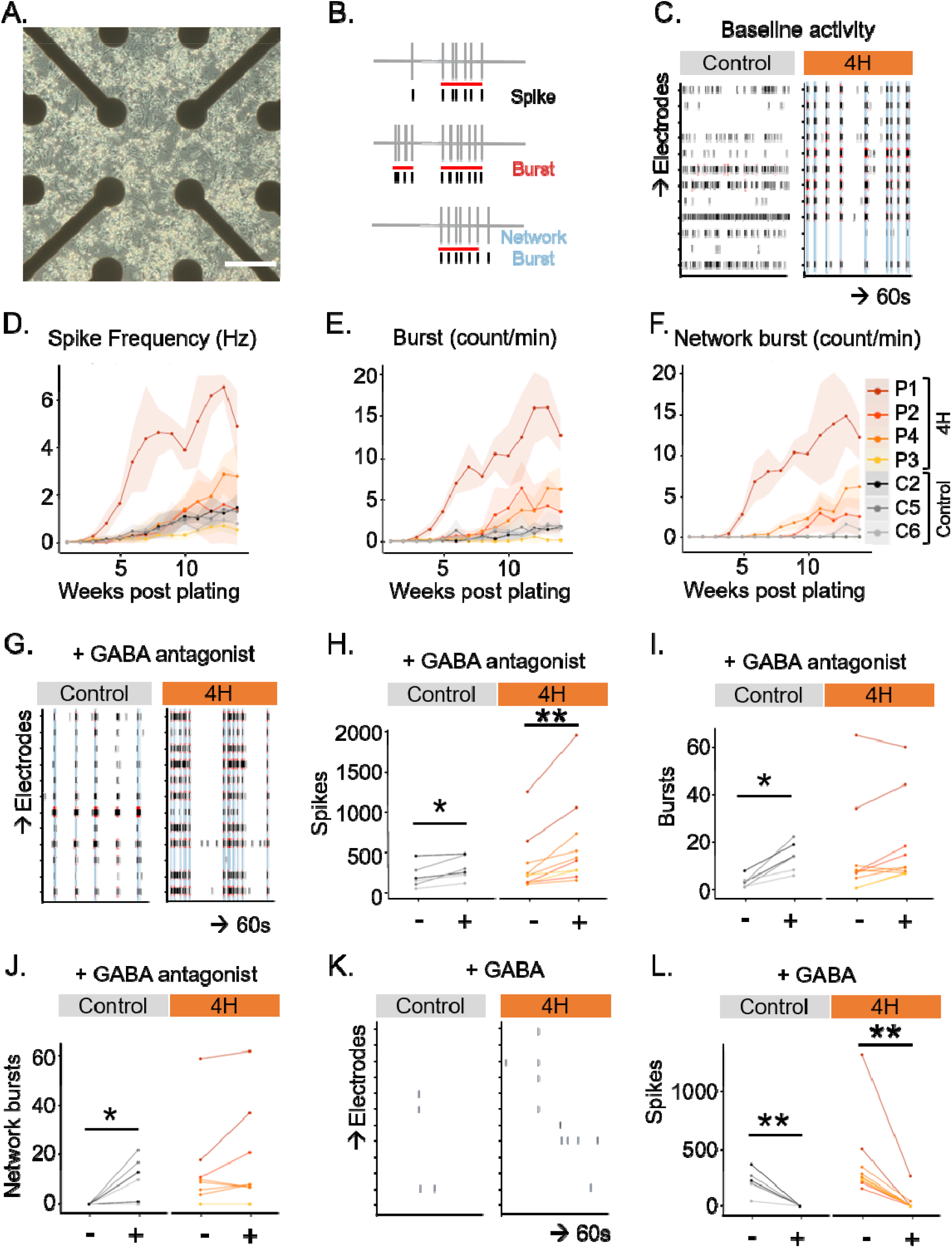
4H neurons show high network activity and a reduced response to GABA antagonists. (**A**) shows a brightfield image of neurons plated on MEA. (**B**) shows schematic example of spike and burst detection from MEA recording. (**C**) shows a representative example of baseline activity at 14 weeks post plating over 60 seconds on different electrodes in control and 4H neuronal cultures. Of the 4H neurons 3 out of 4 show an increased spike frequency (**D**), burst count (**E**) and network burst count (**F**) than control lines, although the differences are not statistically significant. (**G**-**J**) After addition of GABA antagonists bicuculline and gabazine, control neurons show a significantly increase in spikes (**H**), burst (**I**) and network bursts (**J**), while 4H neurons only show an increase in spikes (**I**). (**K**-**L**) Addition of GABA decreased the number of spikes in both control and 4H neurons. Scale bar = 200 μm, Hz = hertz. * = P < 0.05, ** = P < 0.01 (**D**-**F**) data points and lines represent averaged data per patient, with the light areas representing SD per patient. (**H, I, J, L**) data points and lines represent data per well, with lines with the same color representing wells from the same individual.

To establish whether increased network activity in 4H neuronal networks is due to altered synaptic signaling, modulators for glutamate or GABA signaling were added to the cultures at 14 weeks post plating. Network activity was measured for 10 min before and after addition of modulators. Ionotropic AMPA receptor antagonist DNQX (10 uM) or NMDA antagonist APV (50 µM) were used to measure glutamatergic signaling. The inhibition of either one of these glutamate receptors resulted in a significant decrease of spike count for both 4H and control networks (control cells DNQX before 398.1 ± 111.5 after 221.3 ± 103.0, *Z* = −2.201, *P* = 0.028; control cells APV before 174.6 ± 121.7 after 107.7 ± 109.2, *t*(5) = 5.369, *P* = 0.003; 4H cells DNQX before 834.3 ± 424.1, after 126.6 ± 65.1, *t*(9) = 3.412, *P* = 0.008, 4H cells APV before 174.6 ± 121.7, after 107.7 ± 109.2, *Z* = −2.803, *P* = 0.005; Supplementary Fig. 1B-C,E-F). In line with the spike counts, burst and network bursts mostly disappeared (data not shown), showing that coordinated network activity depended on glutamatergic signaling and showed no obvious differences between 4H and control lines.

To study GABAergic signaling, GABA_A_ receptor antagonists Bicuculline (30 µM) combined with Gabazine (20 µM) or the neurotransmitter GABA (10 µM) were administered. After addition, GABA antagonists evoked network bursts in controls that were mostly absent before (burst before 2.85 ± 2.00, after 13.5 ± 5.00, *Z* = −2.201, *P* = 0.028; network burst before 0 ± 0, after 10.5 ± 6.99, *Z* = −2.023, *P* = 0.043, Fig. 5G-J). In contrast, the number of bursts and network bursts did not significantly change in 4H patients (burst before 14.13 ± 12.57, after 18.06 ± 11.60, *Z* = −1.886, *P* = 0.059; network burst before 11.7 ± 10.91, after 15.00 ± 12.45, *Z* = −1.527, *P* = 0.127, Fig. 5G-J). Administration of GABA decreased neuronal activity in both control and 4H cells (control before 215.13 ± 83.50, after 1.68 ± 0.25, *t*(5) = 5.005, *P* = 0.004; 4H before 374.73 ± 212.51, after 32.18 ± 50.59, *Z* = −2.803, *P* = 0.005, Fig. 5K,L), showing normal post-synaptic GABAergic response.

This data shows that the decreased generation of GABAergic synapses correlates to an increase in network activity, and there is a decreased response of 4H neurons to treatment with GABA antagonists.

### 4H neurons show robust myelination *in vitro*

A subset of human interneurons, mainly the parvalbumin neurons, is myelinated(35) and one of the major characteristics of 4H is hypomyelination. Therefore, we decided to analyze oligodendrocyte maturation and myelination on human neuron-oligodendrocyte co-cultures. We altered a previously published protocol for neuron-OPC culture(23) to generate a myelinating co-culture. Indeed, after 40 days of co-culture we observed a robust generation of mature oligodendrocytes (defined as MBP^+^ cells) and myelination, shown by overlap between MBP and NF200 staining (Fig. 6A-B). The presence of myelin was confirmed by electron microscopy, which showed compacted myelin sheets around neurites (Fig. 6C).

**Figure 6.**
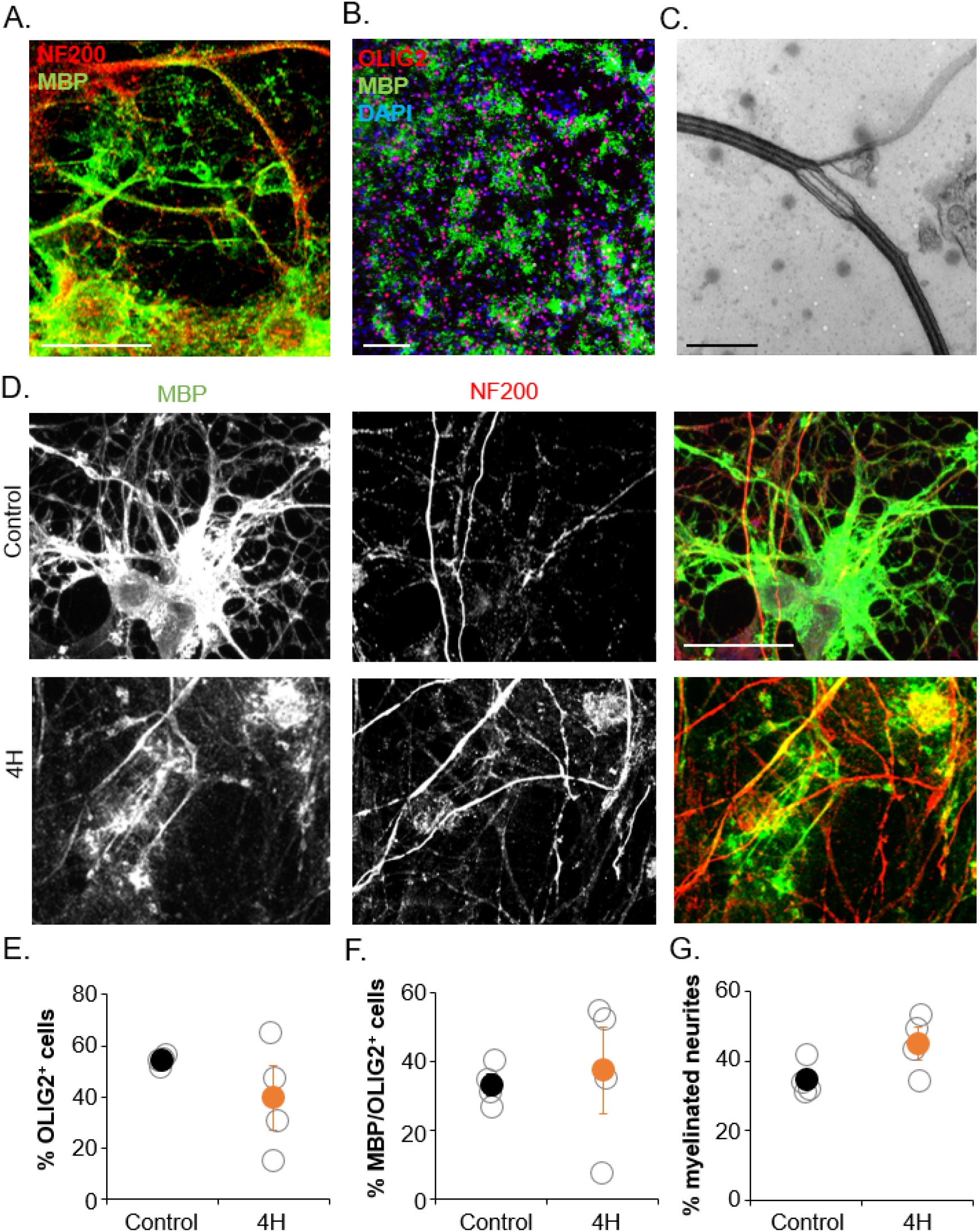
Oligodendrocyte maturation and myelination in co-culture with 4H neurons is normal. (**A-B**) shows that neuron-oligodendrocyte co-cultures show robust generation of oligodendrocytes and myelination. (**A**) Staining for myelin protein MBP and axonal marker NF200 shows wrapping of oligodendrocyte processes around axons. (**B**) Staining for MBP and early oligodendrocyte marker OLIG2 shows a robust generation and maturation of oligodendrocytes. (**C**) Electron microscopy confirms the presence of compact myelin sheets around neuronal processes. (**D**) Cultures with 4H or control neurons and control oligodendrocytes both show oligodendrocyte maturation and myelination. Quantification showed no changes in oligodendrocyte generation, defined as the percentage of DAPI^+^ cells that were OLIG2^+^ (**E**). Similarly, oligodendrocyte maturation (**F**; percentage of OLIG2^+^ cells that developed in MBP^+^ cells) or the percentage of neurites that is myelinated (**G**) was not changed in cultures with 4H neurons. (**A**,**D**) Scalebar = 25 μm, (**B**) scalebar = 100 μm, (**C**) scalebar = 500 nm. (**E**-**G**) open circles represent average per individual patient/control, solid circles represent the mean per genotype. Error bars represent standard error of the mean.

Next, the myelinating cultures were applied to 4H cell lines. Cortical neurons of 4H or control patients were co-cultured with control glial cells. After 40 days of co-culture cells were fixed and stained for neuron (NF200) and oligodendrocyte markers (OLIG2 and MBP). In both 4H and control cultures oligodendrocytes matured into MBP^+^ cells and showed myelination of neurites (Fig. 6D). On average there was no statistical significant change in the generation of oligodendrocyte lineage cells in 4H cultures, although 3 out of 4 patient lines showed a decrease in the percentage of OLIG2-positive cells compared to controls (control 54.3 ± 1.86; 4H 39.6 ± 21.10, Fig. 6E). In 4H cultures, the percentage of mature oligodendrocytes (MBP/OLIG2^+^ cells) was about 35% for cultures with control and 4H lines, but the variation between 4H lines was high (37.5 ± 21.36), while maturation was quite stable between control lines (33.3 ± 5.59, Fig. 6F). Myelination was analyzed by measuring the percentage of NF200^+^ neurites that show co-localization with MBP staining. No changes in the percentage of myelinated neurites were observed between cultures with 4H and control neurons (control 34.7 ± 4.85; 4H 45.1 ± 8.05, Fig. 6G). While this neuron-glia culture set-up provides a novel tool to study myelination defects in brain diseases, we could not identify changes in 4H. This might be challenging if e.g. only a specific neuronal subtype shows affected myelination.

### Targeting the SHH pathway does not improve 4H interneuron phenotypes

ARX is an important regulator of SHH gradients during development(36), suggesting that the decreased expression of *ARX* in 4H might affect interneuron development through the SHH pathway. We tested the prospects for targeting the SHH pathway in 4H in the cortical neuronal cultures. In a pilot study it was established that twice a week treatment with SHH pathway agonist SAG at 100 nM from day 18 onwards increased the percentage of GABAergic synapses on a 4H line, without any (negative) effects on a control line (Supplementary Fig. 2A). PCR analysis confirmed upregulation of SHH target *GLI1* in 4H cells after treatment with 100 nM SAG for 2 weeks (Supplementary Fig. 2B-C). As such, all 4H and iPSC lines were treated with 100 nM SAG from day 18 until the end point of the experiment at day 56. Morphology analysis of neurons showed that SAG treatment decreased the number of dendritic segments, extremities and dendritic length per cell in control cells, suggesting SAG treatment may affect the dendritic complexity (segments vehicle 6.34 ± 1.07, SAG 4.29 ± 1.12 *t*(6) = 2.481, *P* = 0.024; extremities vehicle 3.34 ± 0.39, SAG 2.36 ± 0.46, *t*(6) = 3.176, *P* = 0.010; length vehicle 283 ± 25, SAG 193 ± 17, *t*(6) = 3.194, *P* = 0.019; Fig. 7A-B). In 4H cells, no significant effect of SAG treatment on neuronal morphology was observed. A paired samples test comparing vehicle and SAG treatment for all neuronal batches showed that SAG treatment significantly increased the percentage of VGAT^+^ synapses compared to vehicle treatment (vehicle 31.3 ± 10.19, SAG 34.6 ± 9.93, *t*(20) = − 2.125, *P* = 0.046, Fig. 7C-D). However, the increase was small and no significant improvement in the 4H lines specifically was observed (Fig. 7C-D). Although SAG treatment was able to increase expression of SHH target *GLI1*, it was not able to improve 4H interneuron development *in vitro*.

**Figure 7.**
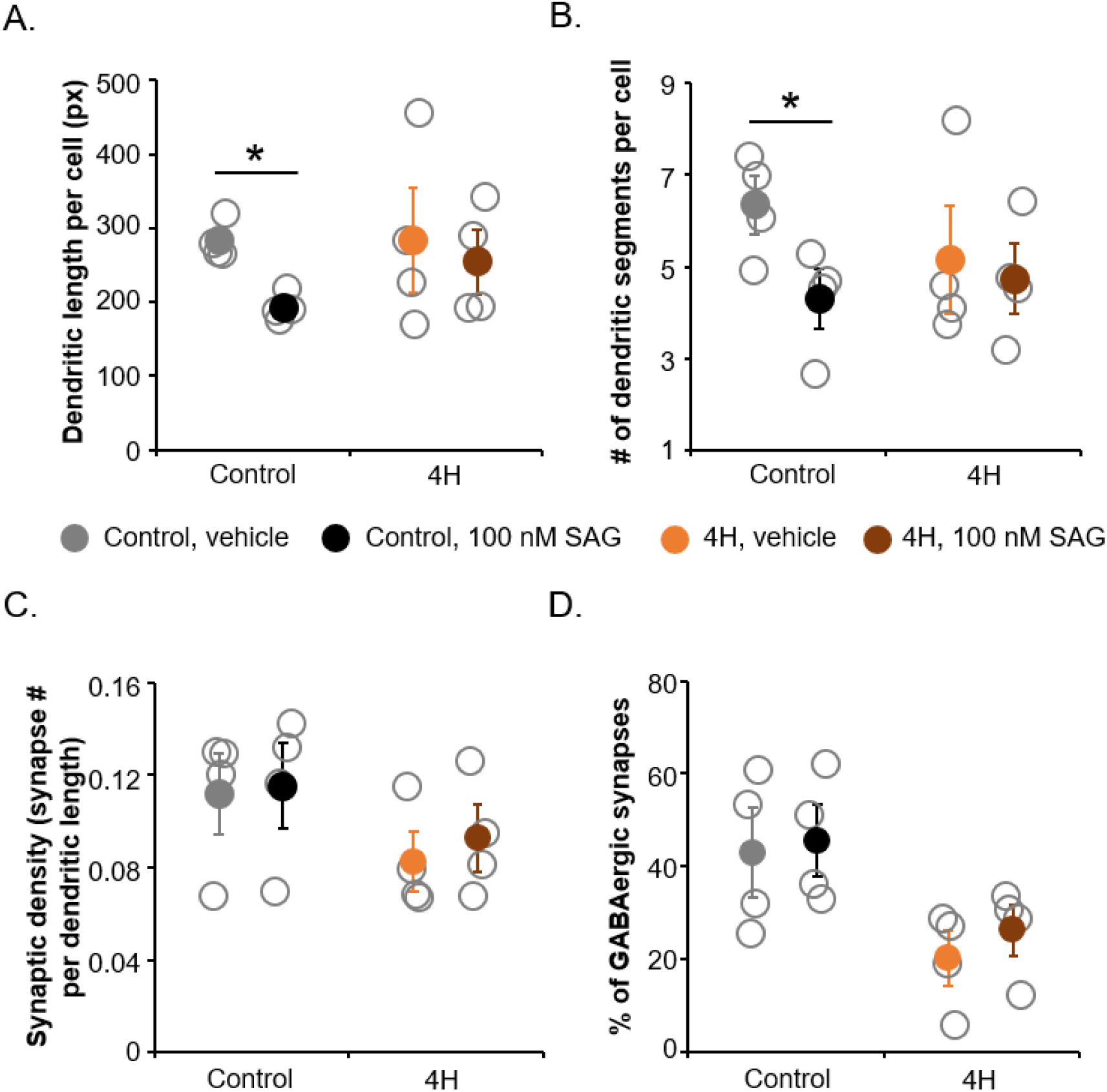
Targeting the SHH pathway in cortical neurons did not improve 4H associated GABAergic phenotypes. Control and 4H neurons were treated with vehicle (DMSO) or 100 nM SAG from day 18 to day 56. After SAG treatment, the dendritic length per cell (**A**) and number of dendritic segments per cell (**B**), both based on MAP2 staining, were significantly decreased in control but not in 4H neurons. No significant changes in the number of synapses per dendritic length (**C**) or the percentage of GABAergic synapses (**D**) were observed. Px = pixels. * = P < 0.05 Open circles represent average per individual patient/control, solid circles represent the mean per genotype. Error bars represent standard error of the mean.

### 4H mutations affect parvalbumin interneuron lineage

Next, we aimed to identify whether specific cortical interneuron subtypes are affected in 4H. Primers targeted the five major human interneuron subtypes: somatostatin (SST) neurons, parvalbumin (PV) neurons, vasoactive intestinal peptide (VIP) neurons, inhibitor of DNA binding 2 (ID2) neurons and neuron derived neurotrophic factor (NDNF) neurons(37). The expression of NEUN was decreased in 4H cultures at day 30, although the difference was not significant (control 1.27 ± 0.82, 4H 0.41 ± 0.25, *t*(8) = 2.171, *P* = 0.062, Fig. 8A). However, since a difference in the number of neurons may affect the relative expression of GABAergic markers when only corrected for a housekeeping gene, all other markers were corrected for expression of both *EIF4G2* and *NEUN. DLX2* is important for development of cortical interneurons and downregulated upon maturation of neurons(38). Indeed, in control cells the *DLX2* expression decreased between day 30 and day 56, which was less pronounced in 4H cells (control day 30 1.98 ± 1.90, day 56 0.31 ± 0.12; 4H day 30 2.06 ± 1.01, day 56 1.61 ± 1.70; Fig. 8B). Despite the decreased number of GABAergic synapses in 4H cultures, markers for specific interneuron subtypes were not significantly decreased in 4H cultures (Fig. 8C-H). In contrast, the expression of *ERBB4*, a marker for PV neurons, was significantly increased in 4H neurons compared to controls (day 30 control 2.28 ± 3.47, 4H 21.45 ± 19.51, *Z* = −2.132, *P* = 0.038; day 56 control 0.42 ± 0.09, 4H 2.50 ± 2.76, *t*(6) = −2.710, *P* = 0.035, Fig. 8H). This suggests that the generation of specific interneuron subtypes is not impaired in 4H, but rather the development or maturation of interneurons may be affected.

**Figure 8.**
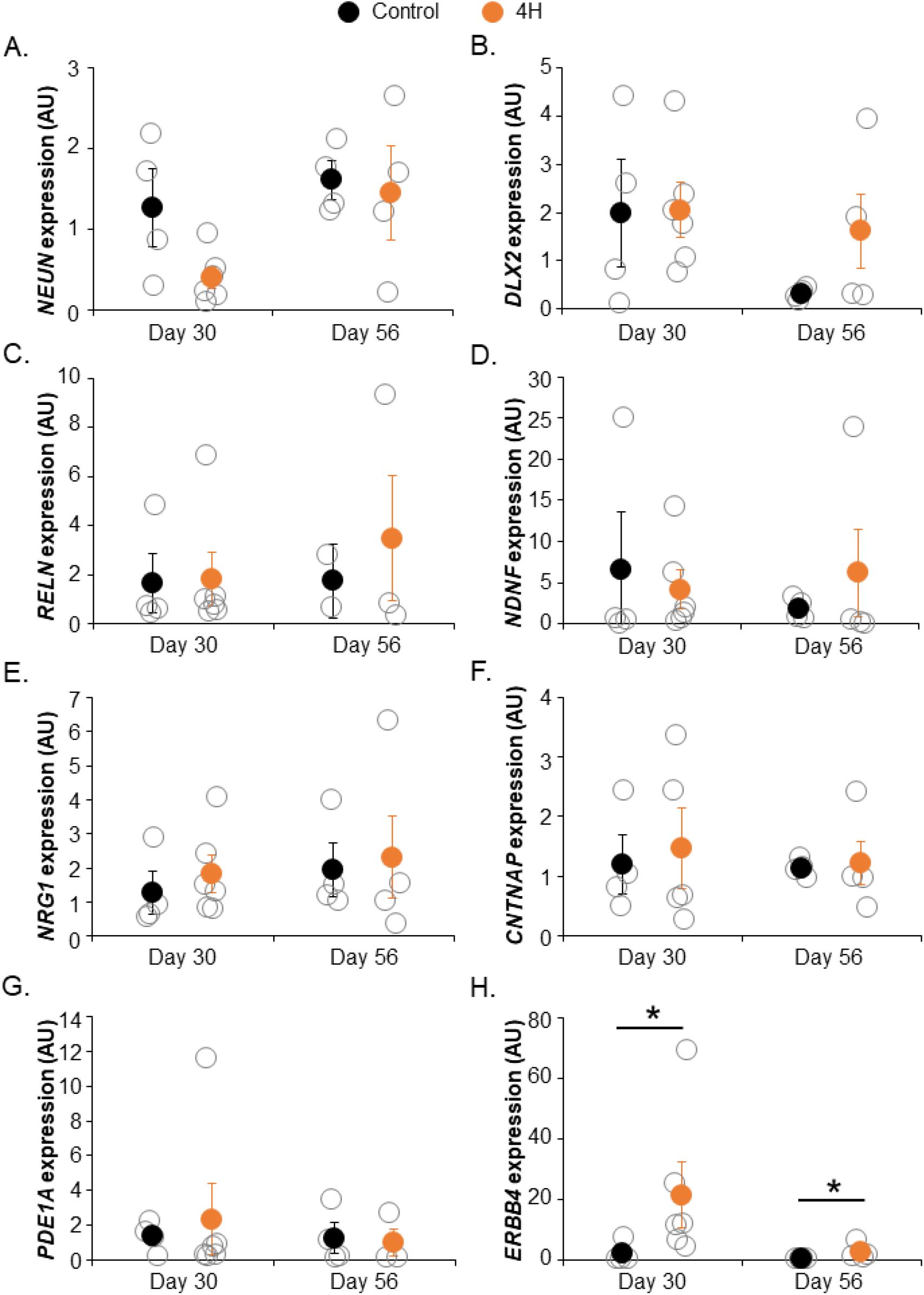
4H neurons show increased expression of parvalbumin interneuron marker ERBB4. QPCR analysis for neuronal and GABAergic markers on day 30 and day 56 control and 4H neurons. (**A**) shows NEUN expression, which is decreased in day 30 4H neurons, therefore (**B**-**H**) are corrected for both NEUN and EIF4G2 expression. (**B**) Expression of DLX2 decreased over time in control neurons, which is less pronounced in 4H cells. (**C**-**H**) show qPCR analysis of specific interneuron subtypes: (**C**) RELN, an early marker for NDNF interneurons, (**D**) NDNF, (**E**) NRG1, an early marker for ID2 interneurons, (**F**) CNTNAP, an early marker for VIP interneurons, (**G**) PDE1A, an early marker for SST interneurons and (**H**) ERBB4, an early marker for PV interneurons. ERBB4 expression was significantly increased in 4H neurons at day 30 and day 56 (**H**). AU = arbitrary units, * = P < 0.05

## Discussion

Although the genetic defects causing 4H leukodystrophy have been identified, our knowledge of underlying molecular mechanisms and the affected cellular subtypes is lacking. This study aimed to get more insight into the affected brain cell types and pathways. We started with an unbiased RNA sequencing screen on 4H and control fibroblasts, iPSCs and cerebellar cells. Although only a small number of genes were differentially regulated in all cell types, an interesting finding was the decreased expression of *ARX* in cerebellar cells of 4H patients. Considering the important role of ARX in cortical neuronal development and migration(32, 33, 39), we studied GAD65/67 expression in patient tissue and showed an increase in GAD65/67 immunoreactivity confirming interneuron changes in the cortex of 4H patients. In iPSC-derived cortical neuron cultures, 4H cortical neurons also showed decreased expression of *ARX* and affected interneuron generation as measured by a decreased percentage of GABAergic synapses. The altered synaptic ratio has functional consequences, as it was correlated to an increased network activity. A decreased GABAergic signaling in 4H also became apparent after treatment with GABA antagonists bicuculline and gabazine. 4H neurons did not show changes in activity upon treatment with the GABA antagonists, while control cells showed a significantly increased activity after treatment. As ARX was reported to have important roles in the Parvalbumin interneurons(39), we tested whether specific interneuron lineages were affected in 4H. Indeed QPCR analysis identified an increase in *ERBB4*, a marker for PV neurons, confirming other results that interneuron regulation may be affected in 4H. Together, interneurons are affected in 4H and we show decreased ARX expression in different neuronal subtypes using iPSC models. Cortical neuron cultures identified a decreased number of inhibitory synapses, increased network activity, and the results are specifically pointing to defects in the PV neurons.

We focused on the involvement of ARX in 4H leukodystrophy because of its role in cortical development and it association with other pleiotropic disorders. Loss of function mutations in ARX lead to pleiotropic disorders such as X-linked Lissencephaly with Ambigious Genitalia (XLAG, OMIM 300215), Agenesis of the Corpus Callosum with Abnormal Genitalia (OMIM 30004), and Lissencephaly with cerebellar hypoplasia(19). ARX impacts interneuron generation, development and migration(40) and loss of ARX expression in interneurons alters their excitability and causes epilepsy in mice(33, 39). In animal models it was shown that Arx acts with FoxA2 to regulate expression of Shh(36). SHH plays important roles in the development of granule cells (and thereby cerebellar volume); oligodendrocytes (and thereby myelinating cells and myelination); jaw and teeth; and the eyes(41-47). It is possible that POLR3 disorders, and 4H in particular, involve ARX-related pathway defects, where altered *ARX* expression (rather than gene mutation) works to drive clinical phenotype through an ARX to SHH pathway.

Although downregulation of ARX may have significant impact on neuronal development, we could not confirm the SHH pathway as a therapeutic target in 4H in our disease model. Neurons were treated with SAG starting from day 18 to day 56 of differentiation. At day 18, the neuronal batches undergo quality control, before plating for generating mature neuronal cultures or frozen for later use. It is possible that treatment starting from day 18 is too late and some of the developmental alterations following a decreased expression of *ARX* and a dysregulated SHH pathway, have already taken place. However, this would mean that targeting the SHH pathway is not a viable option for patients, who are generally diagnosed postnatally. Another explanation may be that the SAG concentration was not high enough to activate the SHH pathway, although, the expression of SHH target *GLI1* was increased after SAG treatment. Finally, it is possible that the affected development of cortical interneurons is caused either by mechanisms that do not involve ARX, or via a SHH-independent pathway. For example, ARX may affect histone demethylation through KDM5C(48, 49) which has been implicated in neurodevelopmental disorders(50). In conclusion, SAG treatment did not revert the neuronal phenotype in our cortical 4H cultures, and following studies should reveal whether a changed treatment protocol or different target would have more beneficial effects.

The percentage of GABAergic synapses was significantly decreased in 4H cultures, suggesting an alteration in the synaptic balance. Indeed, network activity measurements showed a higher activity in most 4H cultures, although results did not reach statistical significance. Interestingly, there was a significant correlation between the percentage of GABAergic synapses and the amount of bursts and network burst, showing that synaptic balance changes also have consequences for functional network behavior. This was also confirmed by the measurement of network activity after treatment with GABA antagonists. In control cultures, the network activity significantly increased after bicuculline/gabazine treatment that releases synaptic inhibition by blocking GABA-A receptors. However, in 4H cultures treatment with GABA antagonists did not significantly change network activity, suggesting that there was less GABAergic signaling that could respond to antagonistic inhibition. Interestingly, the activity in 4H cultures did decrease after addition of GABA, which suggests a normal post-synaptic GABAergic response in 4H cells. The decreased number of inhibitory synapses could be caused by several mechanisms, e.g. reduced network maturation, a reduction in the total or a specific subset of interneurons, less (active) inhibitory synapses per neuron, or an increase in the number excitatory synapses. Follow-up studies should give more insight into this. Also, as patient lines present differential changes, possibly in line with the broad clinical presentation, an increased panel of patient lines would be advised that could possibly also support personalized medicine development.

In light of our *in vitro* findings, it seems contradictory that GAD65/67 immunoreactivity is higher in 4H post mortem tissue. It could possibly be explained if only a subpopulation of interneurons is affected in 4H, as it is reported that interneuron subtypes have different levels of GAD65(51) or it can be indicative of axonal reorganization of remaining interneurons.(52) However it is difficult to determine if that is the case because we co-stained for GAD65 and GAD67 isoforms. Alternatively, the increased GAD65/67 expression does not reflect changes in the interneuron populations, but is rather the consequence of hyperexcitability. Epilepsy is a common feature in 4H and human post mortem studies on temporal lobe epilepsy have also identified increased levels of GAD67, combined making this a plausible explanation.(53) Also described in models of epilepsy, upon neuronal excitotoxicity and increased glutamate release, GAD65/67 is upregulated as compensatory mechanism to convert excess of glutamate into GABA(54). However, while postmortem analysis confirmed network changes in 4H, follow-up studies are warranted, although complicated by lack of human primary tissue.

Alterations in GABAergic activity may underlie 4H associated neurological signs like epilepsy, but also hypomyelination. It is currently unknown how POLR3 mutations cause hypomyelination. Coulombe et al.(55) hypothesized that hypomyelination in 4H is either caused by a currently unknown POLR3 target that plays a key role in myelin biogenesis, or that POLR3 mutations lead to a globally reduced transcription at a crucial milestone in oligodendrocyte development. We investigated an alternative hypothesis, where the altered myelination is secondary to neuronal dysfunction. Cortical (parvalbumin) interneurons in the human brain are myelinated(35, 56), and myelination can be regulated by GABA receptor activity on oligodendrocytes(57). To study whether the decreased interneuron generation in 4H cortical cultures also affected myelination, a new co-culture protocol was developed. This co-culture of human neurons and glial cells show robust maturation of oligodendrocytes and myelination of human axons, a challenging phenomenon to study *in vitro*. No differences in oligodendrocyte maturation or myelination were observed between cultures with control or 4H neurons. This could suggest that the hypomyelination observed in 4H patients is caused by an oligodendrocyte intrinsic effect, rather than mediated through affected neurons. However, it is also possible that our culture set-up was not sensitive enough to pick up changes in myelination for 4H. The neurons used for the co-culture consist of a mix of GABAergic and glutamatergic neurons, so a myelination defect on interneurons may be masked by normal myelination of glutamatergic neurons. Additionally, PV neurons mature during late development and not all neurons in the presented co-culture system showed myelinated processes. Although the novel neuron-glia co-culture can measure myelination *in vitro*, further improvement of these model systems may be necessary to identify mechanisms underlying hypomyelination in 4H leukodystrophy.

We aimed to identify the interneuron subtype that was most affected in 4H cultures. To our surprise, we were not able to identify a significant decrease in any of the five major cortical interneurons populations(37). Instead, we found a significantly increased expression of *ERBB4*, an early marker for PV neurons. There are a few possible explanations for the discrepancy between the decrease in GABAergic synapses and the increase in *ERBB4* expression. It is possible that interneuron generation in 4H is normal, but that interneurons are affected during later development or maturation stages. As *ERBB4* is an early marker for PV neurons, it is also possible that the increase reflects a lack of maturation, rather than an actual increase in PV neurons. A defect in maturation would be consistent with the *DLX2* expression, which decreased in control cells between day 30 and day 56, but not in 4H cells. Unfortunately, PV neurons mature late in cortical development (late gestation/postnatal(58-60)), and we were not able to measure *PVALB* expression in our neuronal cultures. PV interneurons are the largest group of GABAergic neurons in the cortex. They are fast spiking interneurons with a high metabolic demand, which makes them vulnerable for injury(61). Interestingly, the development of SST and PV neurons is depending on SHH levels(62, 63), and deletion of *Arx* in PV neurons specifically led to increased neuronal activity and altered synaptic properties in a mouse model(39). Overall, our data suggests that PV interneurons may be specifically affected in 4H and should be further studied.

To conclude, transcriptome analysis of cerebellar cells differentiated from patient iPSCs revealed ARX as a potentially important regulator of 4H brain pathomechanisms. Indeed, a decreased expression of *ARX* was confirmed in cortical neuron cultures, which showed affected generation of GABAergic synapses consistent with known effects of ARX on interneuron development. The decrease in GABAergic synapses correlated to increased neuronal network activity. QPCR analysis revealed alterations in *ERBB4* expression, suggesting the PV interneurons might be of particular interest for 4H. Although more research is needed, this study provides a first insight into specific brain cell types and pathomechanisms affected in 4H leukodystrophy.

## Abbreviations

APV: (2R)-amino-5-phosphonovaleric acid
ARX: aristaless related homeobox
DEG: differentially expressed gene
DNQX: 6,7-dinitroquinoxaline-2,3-dione
ID2: inhibitor of DNA binding 2
iPSC: induced pluripotent stem cell
NDNF: neuron derived neurotrophic factor
NMM: neural maintenance medium
MEA: multi electrode array
POLR3: RNA polymerase
III PV: parvalbumin
t-SNE: t-distributed stochastic neighbor embedding
TTX: tetrodotoxine
SHH: sonic hedgehog
SST: somatostatin
VIP: vasoactive intestinal peptide

## Acknowledgements

The authors would like to thank Kyoko Watanabe and Danielle Posthuma for analyzing the RNA sequencing data, Jan van Weering, Joke Wortel and Rien Dekker for assistance with electron microscopy, and Brendan Lodder for assistance with RNA isolation. NIW is member of the European Reference Network for Rare Neurological Disorders (ERN-RND), project ID 739510.

## Author contributions

SD designed and executed experiments with cortical neurons. LK executed MEA experiments and qPCR analysis on cortical neurons together with SD and NB. DBH designed and executed

RNA sequencing experiments. MB and MB processed and analyzed human postmortem tissue. NIW and VMH supervised the project. SD, LK, NIW and VMH wrote the manuscript with valuable contributions of all authors.

## Supplementary material

Supplementary material is available online.

