## Supplementary Methods and Figures for "Cortical interneuron development is affected in leukodystrophy 4H"

**RNA isolation**

Total RNA was isolated from samples in Trizol using a chloroform-isopropanol extraction procedure. Briefly, cells were washed 3x with PBS (Braun Medical) and then incubated in 750uL Trizol for < 2 min at room temperature (RT). Samples in Trizol were transferred to 1.5mL Eppendorf tubes (VWR), dissolving any clumps with gentle titration, and stored at -80 C until needed for RNA isolation. After thawing sample tubes at RT, 150uL of chloroform was added to samples in Trizol, solutions mixed by shaking, and tube allowed to stand for 10min at RT. Tubes were then centrifuged (>13K g, 10min, 4C) and the top, clear liquid layer containing RNA transferred to new tubes. RNA was then precipitated by addition of 350uL isopropanol (Sigma-Aldrich) for every 750uL Trizol in the original sample. The tubes were inverted gently to mix, and the solution was allowed to stand for 10min at RT. The samples were centrifuged again (>13K g, 10min, 4C) and the supernatant was removed from the pellet. The pellet was then washed 2x with EtOH (VWR), with centrifugation after each wash step (7500g, 5min, 4C). The pellet was allowed to air-dry before dissolving in appropriate quantity of DMPC water and stored at -80C until needed. RNA concentration and quality were initially measured on a Nanodrop 3000.

**RNA sequencing**

Prior to RNAseq, concentration and quality were confirmed using RNA Analysis ScreenTape (Agilent Technologies, USA) on a 2200 TapeStation System (Agilent) to establish RIN scores. RNA samples of sufficient quality were processed according to manufacturer’s instructions using the TruSeq Stranded Total RNA Library Prep Kit with Ribo-Zero Human (Illumina Inc., USA), generating tagged cDNA libraries capable for high-throughput RNA sequencing. Library concentration and quality were confirmed using D1000 ScreenTape (Agilent Technologies, USA) on the 2200 TapeStation System (Agilent). Samples were run on a Illumina HiSeq2500 at SR50.

Quality control was performed using FastQC, and sequencing reads were aligned to Human Genome hg38 using Bowtie2(1) with default parameters. Aligned reads were converted into count per gene using featureCount function from Rsubread package(2) with gene annotations obtained from GENCODE v26 (https://www.gencodegenes.org/) which contains unique 63,199 genes. Reads Per Kilobase per Million (RPKM) were further computed by using rpkm function from edgeR package(3). Genes were filtered on such with count per million (CPM) > 1 in ≥ 50% of samples per cell type which resulted in 21,542 unique genes including non-coding RNAs.

**Sample t-SNE map**

To evaluate similarity between samples, we applied the t-Distributed Stochastic Neighborhood Embedding (t-SNE) to the RNA-seq expression profiles. The t-SNE non-linearly projects local similarities between samples at the cost of retaining the similarities between dissimilar samples. We first normalized RPKM; zero-mean normalization followed by log2 transformation with pseudo count 1. Then t-SNE was performed 100 times and we selected the solution with the lowest Kullback-Leibler divergence.

**Cell type validation with ENCODE samples**

Gene expression profiles of 2 iPSC samples and 2 granule cell samples were obtained from ENCODE (ENCSR722POQ for iPSCs and ENCSR313IUO for granule cells).(4) For expression profiles of ENCSR722PQO, obtained read counts were converted into RPKM as described above. For ENCSR313IUO, Fragments Per Kilobase Per Milling (FPKM) was available. Genes with expression value zero in all samples were filtered out. Since the expression data is not directly comparable between data sets, we performed Spearman’s rank correlations across these 4 samples from ENCODE and samples from this study by taking intersect genes (18,719 genes in total).

**Differentially expressed gene analysis**

Raw count data was used for differentially expressed gene (DEG) analysis with DESeq2 package.(5) For cell type comparison, we performed DESeq2 for a specific cell type against all other samples and tested all 21,542 genes. Control vs patient DEG was performed per cell type, and genes were further filtered on such with CPM > 1 in ≥ 50% of samples with the testing cell type. The number of tested genes for fibroblast, iPSC and product are 16,890, 18,855 and 19,032 genes, respectively. Genes with absolute log2 fold change (logFC) > 1 and Bonferroni corrected P-value < 0.05 were defined as DEG.

**Cortical neuron differentiation**

Neurons were generated from iPSCs according to previously published protocol.(6) Shortly, iPSCs were differentiated into neural epithelial stem cells (NES) using N2/B27 medium supplemented with Dorsomorphin and SB431. Within 1-2 weeks after plating neural rosettes appeared in the culture, which were selected by manual cutting. NES cells were maintained in N2/B27 medium supplemented with FGF2 and EGF for up to 4 passages. To differentiate NES cells into neurons, medium was changed to N2 medium supplemented with SHH for 4 days, followed by 3 days in Neurobasal/B27 medium with Valproic Acid. The day of the medium change to N2 medium with SHH was considered day 1 of neuronal differentiation. At day 8 of differentiation, cells were split into a new PLO/Laminin coated plate in Neurobasal/B27 medium supplemented with BDNF, GDNF, IGF1 and cAMP. At day 18, Neurons were passaged a final time and either frozen for later use or plated on rat astrocytes for further neuronal maturation. From now on, half of the medium was refreshed twice a week. At day 25, neurons were treated with AraC to remove proliferating progenitors. Neurons were kept in culture until day 56, at which point they resembled mature neurons in morphology and synaptic marker expression.

**Oligodendrocyte differentiation**

Oligodendrocyte progenitor cells (OPCs) were generated from iPSCs according to a previously published protocol(7). Shortly, iPSCs were plated on an anti-adherent plate in N2/B27 medium with EGF, FGF2 and T3 to induce embryoid body formation. From day 2 onwards, RA was additionally added to the medium. At day 10, cells were plated on geltrex coated plates in N2/B27 medium with EGF and T3. At day 18, medium was switched to N2/B27 medium without vitamin A. Throughout the differentiation, cells were passaged 1:2/1:3 into a new well when they reached confluency, using accutase dissociation. Half of the medium was refreshed every other day. At day 37 medium supplements were changed to EGF, FGF2, mouse Laminin, Vitamin C and T3. At day 39, Dorsomorphin was additionally added to the medium. At day 42, medium supplements were changed to mouse laminin, vitamin C, Dorsomorphin and T3. Cells were kept in these medium until day 67 when they were frozen or replated for neuron-OPC co-cultures.

**Supplementary Table 1. Primer sequences**

| **Target** | **Forward primer** | **Reverse primer** |
| --- | --- | --- |
| *ARX* | AAACGCAAACAGAGGCGCTA | CAGTTCCTCCCTGGTGAAGAC |
| *NEUN* | TGGCATGACCCTGTACACAC | GCTGCTGCTTCTCTGTAGGG |
| *EIF4G2* (housekeeping) | AGGACCGCATGTTGGAGATT | TGAGGGGATGGATCCAACTTT |
| *NRG1* | CGTGGAATCAAACGCTACATCT | TTCACCATGAAGCACTCCCC |
| *PDE1A* | CAGCAGTGGACCTGAAGAGTT | TGTGAACTGGTTCTTGCTTCTTG |
| *ERBB4* | GTTCAGGATGTGGACGTTGC | ACACACCGTCCTTGTCAAAGT |
| *NDNF* | GTGCTTAGCATCTTGGCAGG | TGGAGCAGCACCATCCTTAAA |
| *RELN1* | CGTCCTAGTAAGCACTCGCA | TCGCCTAAGTGACCTTCGTC |
| *NEUN* | GTCCCTTACTCCGCCAAGAG | CAAGGTCCTCCTTCTCAGGC |
| *DLX2* | ATCCAGAAATGTGCCTGCGG | AGCATCCTTCCTCCATTGCTT |
| *CNTNAP2* | CGTGGAATCAAACGCTACATCT | TTCACCATGAAGCACTCCCC |

**Immunostaining**

Cells were fixed for immunostaining using either 10 min fixation in 4% PFA (MBP + OLIG2 staining) or by 10 minute fixation in ice cold 1:1 Acetone:Methanol. After fixation, cells were washed 6x 5 min with PBS and incubated in blocking buffer (PBS + 5% NGS + 0.3% Triton X-100 + 0.1% BSA) for 1 hour at RT. Primary antibodies were diluted in blocking buffer and incubated for 1 hour at RT followed by overnight incubation at 4˚C (see Table 2 for primary antibodies). The next day, cells were washed 6x 5min with PBS and incubated in secondary antibodies (Goat-anti-chicken/mouse/guinea pig/rabbit/rat Alexa Fluor 488/568/647) diluted 1:1000 in blocking buffer for 2 hours at RT. Cells were washed 6x 5 min with PBS, incubated in DAPI (1:1000 in PBS) for 1-2 min and embedded on microscope slides using Fluoromount G. Cells were imaged on a Leica DM6000B Fluorescent microscope, or on a Nikon Eclipse Ti confocal microscope for stainings of synapses and myelin.

**Supplementary Table 2. Primary antibodies for immunostaining**

| **Target** | **Host** | **Dilution** | **Company** | **Number** |
| --- | --- | --- | --- | --- |
| MAP2 | Chicken | 1:5000 | Millipore | AB5543 |
| MBP | Rat | 1:500 | Abcam | AB7349 |
| NF-200 | Rabbit | 1:1000 | Sigma | N4142 |
| NF-H | Mouse | 1:100 | Hybridomabank | RT-97-s |
| OLIG2 | Rabbit | 1:500 | Millipore | AB9610 |
| SMI312 | Mouse | 1:1000 | Eurogentech | SMI-312P-050 |
| Synaptophysin1 | Guinea Pig | 1:1000 | Sysy | 101-004 |
| VGAT | Rabbit | 1:500 | Sysy | 131-002 |

**Multi electrode array analysis**

Day 18 cortical neurons, generated as described above, were plated on PLO/Laminin coated MEA plates (Multi Channel Systems 24W300/30G-288) together with rat astrocytes (1:6 ratio) and cultured as described above. MultiChannel Headstage hardware (Multi Channel Systems) and Mulitwell-Screen software (Multi Channel Systems) were used to record baseline electrophysiological activity. Measurements were performed for 30 minutes (after 10 minutes incubation) while controlling the environment at 37°C and humidified 5% CO_2._ Raw data was sampled at 10 kHz and filtered using a high-pass 2^nd^ order Butterworth filter with 100 Hz cut-off and low-pass 4^th^ order Butterworth with 3500 Hz cut-off. The data was analyzed using Multiwell-Analyzer software (Multichannel Systems). Faulty electrodes were manually excluded from the analysis upon visual inspection. Spike detection threshold was set at -5.0 standard deviations from baseline. To start a burst at least 4 consecutive spikes had each to be less than 50 ms apart. Burst events needed a minimal duration of at least 50 ms. If the spikes within a burst are more than 50 ms apart a burst ends, new burst can only start 100 ms after the last burst. Network burst detection was set on 6 (=50%) or more participating channels of which at least 3 (25%) simultaneous. Results were exported and visualized using RStudio, SPSS was used for statistical testing.

To measure the effect of different compounds on neuronal activity in 4H and control cells, cells were treated with either APV (50 µM), DNQX (10 µM), TTX (1 µM), GABA (10 µM) or Bicuculin/Gabazine (30 µM/20 µM), at 14 weeks post plating. In addition, to one well of each line the corresponding solvent (DMSO or H_2_O) was added in the same volume as the compounds. Compounds were mixed in and electrical activity was recorded for 5 minutes at 37°C and 5% CO_2_ flow after a 5 minute incubation period under the same conditions. Plain Neurobasal medium was used to wash out the compounds. After washing, 50% of old and 50% of new Neurobasal/B27 medium with fresh supplements (BDNF, GDNF, IGF, cAMP) were added. Cultures were let to recover for 24h before addition of the next compound.

**ImageJ analysis**

For synaptic analysis the NeuronJ plugin was used to trace MAP2+ dendrites. In SynaptoCount, the dendritic tracing was loaded and used to determine the total number of synapses by counting the number of Synaptophysin1+ puncta present on the dendrites. The amount of VGAT+ synapses was determined by the number of VGAT+/Synaptophysin1+ puncta present on dendrites. The synaptic density was calculated by dividing the total number of synapses by the pixel length of the dendrites. The percentage of VGAT+ synapses was determined by dividing the number of VGAT+ synapses by the total number of synapses.

For myelin analysis the NeuronJ plugin was used to trace NF-200^+^ axons, and the pixel length of axons was measured. Neurite tracings were then loaded on MBP staining, and the pixel length of neurites that showed co-localization of MBP and NF-200 staining was determined. The percentage of myelinated axons was determined by dividing the length of MBP/NF200 double positive neurites by the total length of NF200 neurites.

**Supplementary Figures**

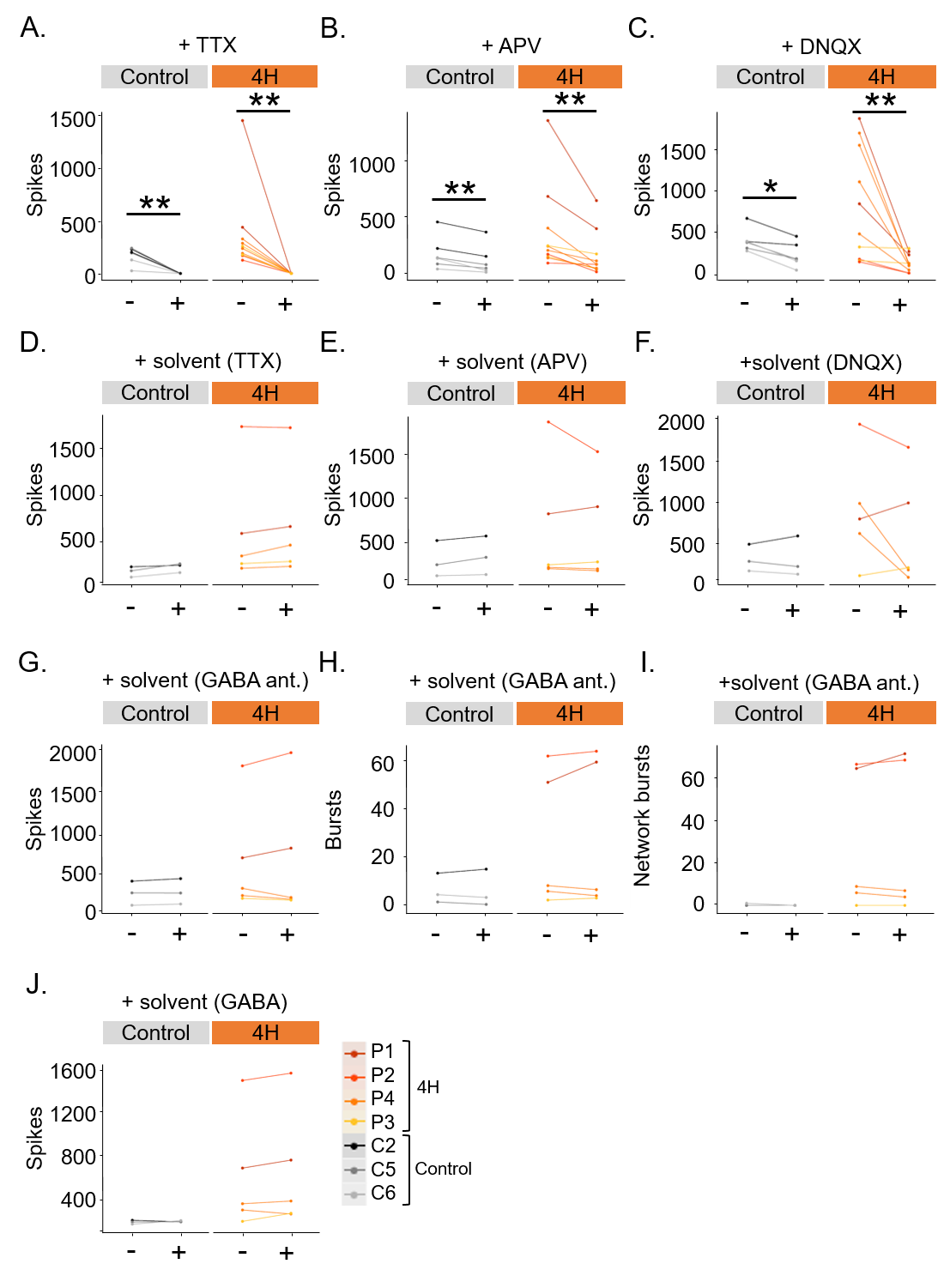

***Supplementary Figure 1. Effect of activity modulators on neuronal cultures.*** *Graphs show that the number of spikes in control and 4H neurons significantly decreased after treatment with TTX (****A****), APV (****B****) and DNQX (****C****). Addition of solvents to cultures did not influence network activity in any of the conditions (****D****-****J****). (****G****-****I****) GABA ant. = GABA antagonists bicuculline and gabazine. * = P < 0.05, ** = P < 0.01. Data points and lines represent data per well, with lines with the same color representing wells from the same individual.*

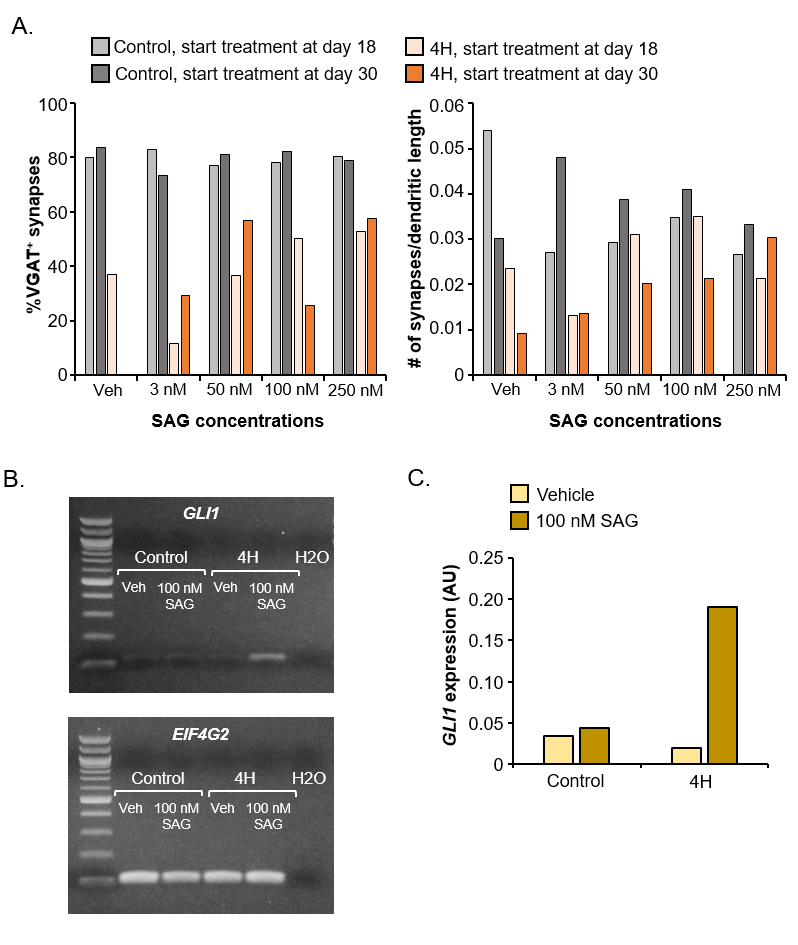

***Supplementary Figure 2. Pilot study to test efficacy of SAG concentrations.*** *A control and a 4H line were treated with vehicle, 3 nM, 50 nM, 100 nM or 250 nM SAG, either started from day 18 or day 30 of the neuronal differentiation. (****A****) The left graph shows the percentage of VGAT^+^ synapses, which did not change in the control line in any treatment condition. However, for the 4H line the percentage of VGAT^+^ synapses was increased after treatment with 100 or 250 nM SAG. The right graph shows the number of synapses per dendritic length. SAG treatment decreased the number of synapses per dendritic length slightly in the control line, although it was slightly increased in the 4H line with 50 or 100 nM SAG when treatment was started at day 18. (****B****) shows the expression of SHH downstream target GLI1 and housekeeping gene EIF4G2 after vehicle or 100 nM SAG treatment in control and 4H cells. Quantification of GLI1 band intensity shows that, after correction for EIF4G2 band intensity, the GLI1 expression is increased in 4H neurons (****C****). Veh = vehicle, AU = arbitrary units.*
